# A Weighted Estimation Approach to Correct for Imperfect Mutation Detection Due to Intra-tumor Heterogeneity in Tumor Sequencing

**DOI:** 10.64898/2026.10.02.756288

**Authors:** Audrey Mauguen, Irina Ostrovnaya

## Abstract

Somatic mutations in sequenced tumors are often detected with imperfect sensitivity. They can be missed because tumor heterogeneity results in some mutations being absent in a sample at hand or present at very low frequencies which, combined with technical limitations like insufficient sequencing depth, reduces their detectability. It becomes apparent with multi-regional sequencing that some mutations confined to specific areas can be overlooked in single-region analyses. Adjusting for such sensitivity becomes critical when mutation frequencies between groups of subjects are compared, such as responders and non-responders to a given therapy. Otherwise, the estimate of the mutation prevalence, or of the difference in prevalences between the groups, can be biased, leading to inflated type I error. Here, we propose a weighted approach to adjust for imperfect sensitivity of the mutation detection and compared it to the unweighted approach. We assume that a surrogate of mutation sensitivity is observed only among patients with called mutation, and we derive its unconditional mean. Such adjustment leads to less biased estimates of mutational frequencies and better control of the type 1 error.

## Introduction

It is understood that somatic mutations occurring during cell development are the driving force of many cancers. However, the impact of these mutations on tumor characteristics, therapy efficacy, or patient outcome is less understood, although widely studied. Many cancer studies now include an estimation of the prevalence of mutations in a given gene and correlate it with certain tumor or patient characteristics. To do so, tumor tissue is collected, either by biopsy or during surgery, DNA is extracted and sequenced, and mutations are called. However, a mutation that is effectively present in the tumor can be missed. This can be due to several reasons. For example, a biopsy only takes a small sample of tumor cells, and since many tumors are heterogeneous, it is possible that the cells containing the mutation of interest are not part of the sequenced sample; or the mutation was not called during the pipeline due to low variant allele frequency.

The chance of observing an existing mutation can be increased by using multiple samples from the same tumor obtained through repeated biopsies, with tissue coming from multiple lesions or regions of the tumor. Although such samples can be insightful, they are rarely obtained and, when they are, it is mostly for research purposes. As an example, multiregional sequencing of kidney tumors performed by Gerlinger et al. (2014) showed that sequencing mutations in a single biopsy may underestimate the true prevalence of the mutation. They estimated TP53 prevalence was 6% using single biopsies of kidney tumors while the prevalence rose to 40% when multiple tumor regions were considered. Similarly, in lung cancer, multiregional whole exome sequencing found a median of *>* 100 additional mutations compared to single sample analysis (Jamal-Hanjani et al. (2017)).

We consider mutations in a context of bulk sequencing of a particular sample from a tumor collected at biopsy or surgery. We assume patients come from two classes, some have a mutation somewhere in the tumor, and others do not. If mutation is called, we assume it is present. If a mutation is not called, the patient could either be among patients without mutations, or among patients with a mutation that was missed. While we are not able to tell these two groups apart on a patient level, we are attempting to adjust the mutation prevalence estimate for those patients that have a mutation but it is missed, using some probability *θ*_*i*_. We are not considering single cell sequencing since it is not often used in clinic, and we focus only on somatic mutations.

Suppose we are testing association between the mutation and binary outcome of interest, defining two groups of patients, cases and controls. If the sensitivity of mutation detection, meaning a chance of detecting a mutation that is present in the tumor, differs between the groups, it can bias the comparison of the mutational prevalences, as illustrated in Figure 1. If the group with the highest mutation prevalence has the lowest sampling sensitivity, then the prevalences might appear similar between the groups, attenuating the correlation between mutation and outcomes. On the other hand, if there is a lower sensitivity in the group with lower mutation prevalence, the difference in prevalences might appear larger than it really is, leading to a false positive association. Thus, mutation detection sensitivity is a function of intra-tumoral heterogeneity which in turn has been shown to be associated with patient outcomes (Morris et al. (2016), Vitale et al. (2021)), since more heterogeneous tumors have more avenues to adapt under therapeutic pressure. Mutations in a gene that could predict favorable patient outcome might be detected with higher sensitivity, potentially obscuring a correlation. It has also been shown that if driver mutations are subclonal, e.g. present only in a subset of tumor cells, they are associated with worse outcome (Landau et al. (2013)). Subclonal mutations are also likely to be present in more heterogeneous tumors and are more likely to be missed, so mutation-outcome association would be confounded by the tumor heterogeneity, resulting in bias.

**Figure 1.**
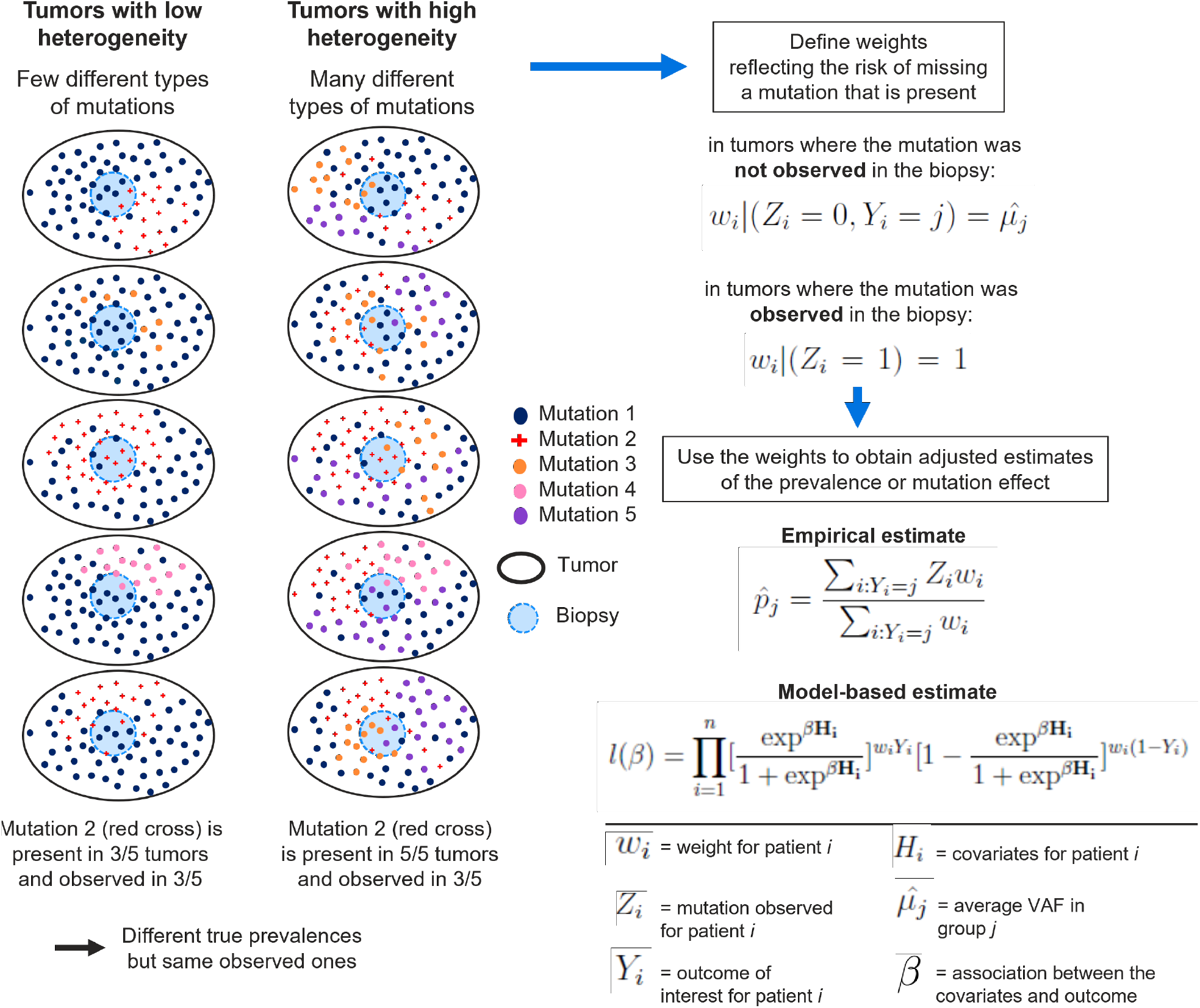
Schema of the study. Illustration of the intra-tumor heterogeneity and summary of the methodological approach.

Previous methodological approaches have been proposed to deal with factors or biomarkers detected with imperfect sensitivity or specificity (ie, subject to misclassification), for example using the EM algorithm (Magder and Hughes (1997)) or when designing clinical trials (Liu et al. (2014)).

## Methods

Suppose *Y*_*i*_ is a binary outcome for patient *i* that takes values *j* = 0, 1. *X*_*i*_ is a binary random variable representing the true mutation status, following a Bernoulli distribution *Ber*(*p*_*j*_) with *p*_*j*_ being the mutation prevalence in group *j*. If the tumor is heterogeneous, *X*_*i*_ might not be observed. Instead, we observe the binary variable *Z*_*i*_ which is equal to 1 if both *X*_*i*_ = 1 and the mutation is observed with probability *θ*_*i*_: *P* (*Z*_*i*_ = 1|*X*_*i*_ = 1) = *θ*_*i*_ and *P* (*Z*_*i*_ = 1|*X*_*i*_ = 0) = 0. Parameter *θ*_*i*_ could be a fixed value *θ*_*i*_ = *θ*_0_ or it could be a random variable with some outcome-dependent distribution *θ*_*i*_|(*Y*_*i*_ = *j*) ~ *f*_*j*_(.), thus the distribution of *θ*_*i*_ can be different by group *j*. If the tumor is homogeneous, *θ*_*i*_ = 1, a present mutation is always detected (we assume that there is no technical error), and the proportion of subjects with an observed mutation is an unbiased estimate of the mutation prevalence. In presence of tumor heterogeneity, *θ*_*i*_ *<* 1, the observed rate in group *j*, denoted as 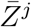, is not an unbiased estimate of the mutation prevalence. Instead, for *i* with *Y*_*i*_ = *j, P* (*Z*_*i*_ = 1|*θ*_*i*_) = *P* (*Z*_*i*_ = 1|*X*_*i*_ = 1, *θ*_*i*_)*P* (*X*_*i*_ = 1) = *θ*_*i*_*p*_*j*_. Unbiased estimate of the mutation prevalence 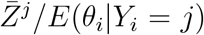. We assume that conditional on outcome group, detection sensitivity is independent of true mutation status.

### Proposed weights

Let’s define weights *w*_*i*_ for each subject. The proposed weights make use of *θ*_*i*_, the probability of observing an existing mutation if it is present in the tumor. An example can be the rescaled variant allele frequency (VAF) used as a surrogate of *θ*_*i*_. In the examples presented below, we rescale VAF using min(1, 2 ∗ *V AF*). However, the VAF is only measured in subjects with an observed mutation.Therefore, while *θ*_*i*_ is needed as the basis for the weights, only the distribution conditional on observing mutation, *f*_*j*_(*θ*|*Z*_*i*_ = 1) can be estimated from the data. The needed unconditional distribution of *θ*_*i*_, *f*_*j*_(*θ*), can be derived as follows.

First, the conditional distribution of *θ*_*i*_ given that the mutation is observed can be written as:

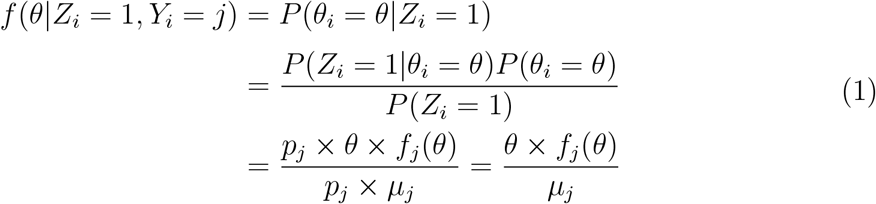

Thus, the distribution of *θ*_*i*_ among subjects with an observed mutation is a size-biased version of its unconditional distribution: mutations with higher detection sensitivity are more likely to be represented among subjects for whom the mutation is observed.

Using Equation (1),

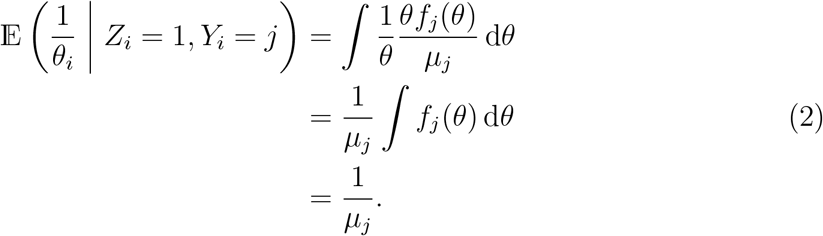

It follows that the unconditional mean detection sensitivity can be expressed as

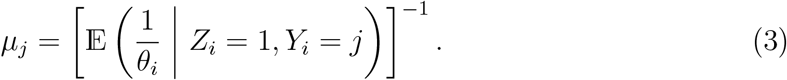

Therefore, *µ*_*j*_ may be estimated from subjects with an observed mutation in group *j* using the sample harmonic mean:

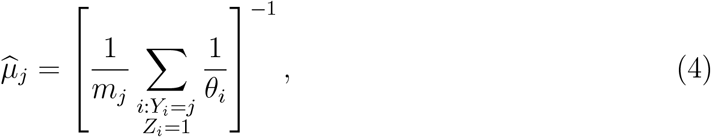

where

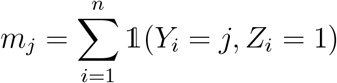

is the number of subjects with an observed mutation in group *j*.

This identity does not require a parametric assumption for *f*_*j*_(*θ*). It does, however, require that *θ*_*i*_ *>* 0 almost surely and that *θ*_*i*_ represent a correctly calibrated probability of detecting a truly present mutation

Let’s define the weights for subjects without observed mutation by 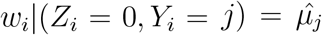, and for the subjects with observed mutation by *w*_*i*_|(*Z*_*i*_ = 1) = 1, while 0 *< w*_*i*_ ≤ 1. Thus, higher value of 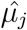 leads to more confidence in true negative status. On the other hand, when 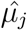 is low, the chances of missed mutation are higher, and *Z*_*i*_ = 0 is downweighted in this case.

Based on these weights, the mutation prevalence in each group can be estimated by

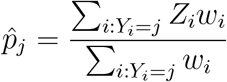

Taking advantage of the definition of weights, the estimated prevalence can be rewritten as follows:

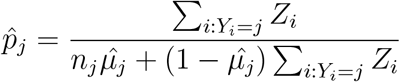

Where *n*_*j*_ = ∑_*i*_ *I*[*Y*_*i*_ = *j*]. Using a Taylor’s expansion, the variance of the estimate can be approximated by

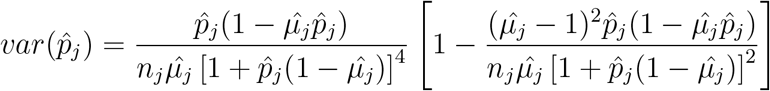

To test whether the difference in mutation prevalences is equal to 0, a Z-test can be used as follows:

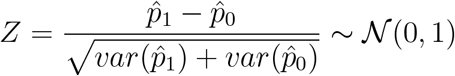

### Weighted logistic regression model

There are studies for which estimating the mutation rate adjusted for other measured characteristics is preferred. An example would be testing association between the response rate to targeted therapy and a given gene mutation, while adjusting for disease stage or mutations on other genes. To do so, a logistic regression model is a common approach used in clinical studies. In order to account for the probability of observing the mutation of interest, the previously defined weights can be used in the regression. As such, observations will be downweighted according to the risk of the mutation being missed.

The likelihood of the weighted model is as follows:

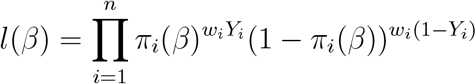

where *Y*_*i*_ is the outcome of interest, *w*_*i*_ is the individual weight as defined above, and *β* measures the strength of association between covariates and the outcome. We define *π*_*i*_(*β*), the individual probability of the outcome, as follows:

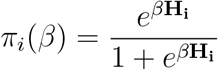

*H*_*i*_ represent the vector of covariates for subject *i*, including the observed mutation status *Z*_*i*_. The parameter estimates are obtained by maximizing the likelihood of the model.

### Simulation Study

#### Simulation Study Settings

A simulation study was conducted to assess the performance of the proposed weighted estimate compared to the unweighted estimate under different scenarios. Two groups *j* = 0, 1 of subjects were simulated. Without loss of generality, they will be called non-responders (*j* = 0) and responders (*j* = 1). The total sample size was fixed at *n* = 400.

In different scenarios we generated patients with different mutation prevalence *p, p* in {0.10, 0.50}, and the odds ratio (OR) of response for mutated vs non-mutated subjects *OR* was 1, 1.5, and 2. The mutation detection sensitivity was defined using a *Beta*(*α*_*j*_, *β*_*j*_) distribution, where the set of parameters {*α*_*j*_, *β*_*j*_} was equal to {10, 20} (corresponding to an average *θ*_*j*_ = 0.33), {15, 15} (*θ*_*j*_ = 0.50), {16, 4} (*θ*_*j*_ = 0.80) or {1, 0} (*θ*_*i*,*j*_ = 1 for all *i* = 1, …, *n*), the latest representing the ideal scenario with no missed mutations.

Each subject was attributed a mutation status using a Bernoulli distribution *x*_*i*_ ~ *Ber*(*p*), where *p* is a mutation rate in the population. The overall probability of response was fixed at *p*_*r*_ = 0.5. Then, logistic regression parameters were defined as *b*_*o*_ = ™*log*((1™ *p*_*r*_)*/p*_*r*_) and *b*_1_ = *log*(*OR*). Each subject was attributed an individual probability of response *p*_*r*,*i*_ based on their mutation status using *p*_*r*,*i*_ = 1*/*{1 + exp[™(*b*_0_ + *b*1 ∗ *x*_*i*_)]}, and a response status *y*_*i*_ ~ *Ber*(*p*_*r*,*i*_). Then, an individual mutation detection sensitivity depending on their response status was simulated using *θ*_*i*,*j*_ ~ *Beta*(*α*_*j*_, *β*_*j*_) from the *Beta* distributions listed above. Thus the presence of an *observed* mutation is simulated using *z*_*i*,*j*_ ~ *Ber*(*p*) × *Ber*(*θ*_*i*,*j*_), where *Ber*(*p*) is the same as used to determine *x*_*i*_.

Once the observations have been simulated, the parameters of interest - regression coefficient *b*_1_, *var*(*b*_1_) and *OR* = exp(*b*_1_) - were estimated using a classical unweighted logistic regression and using the proposed weighted logistic regression, where the covariate entered was the observed mutation status *z*_*i*_ rather than the true mutation status *x*_*i*_. For each scenario, simulations are repeated 1000 times in order to estimate the power and type-I error of the approach. Power is defined as the proportion of simulations with p-value ≤ 0.05 in the scenarios where *OR >* 1, while the type I error is defined as the proportion of simulations with p-value ≤ 0.05 in the scenarios where *OR* = 1. Power and type I error are used to assess the performance of the methods, where the ideal method provides high power while maintaining the type I error below the chosen 5% threshold. In addition, the bias in the estimate of *b*_1_ is assessed. Note the bias is computed relative to log odds ratios between the outcome and the true mutation status, not the observed one.

#### Odds ratio estimates

Results of the simulations are shown in Figure 2. In the scenarios where *θ*_0_ = *θ*_1_, i.e. the number of missed mutations is similar in both groups, both the weighted and unweighted approaches give the same estimates with similar variation. This estimate is unbiased when *θ*_0_ = *θ*_1_ = 1, the scenarios without any missed mutations, as expected, but notice bias (means are far from the black line of true estimate) in most other scenarios (Figure 2). In these scenarios, the weighted approach leads to a lower bias than the unweighted approach. The confidence interval for the weighted estimate contains the true OR in all but three of the 54 scenarios (50% mutation rate; *θ*_1_ = 0.33, *θ*_0_ = 1 and OR = 1.5 and 2; *θ*_1_ = 0.5, *θ*_0_ = 1 and OR = 2), showing acceptable performances. In these three scenarios, the unweighted scenarios showed a far larger bias, and a confidence interval that excludes the true value of the OR, as it does for a total of 21 scenarios out of 54.

**Figure 2.**
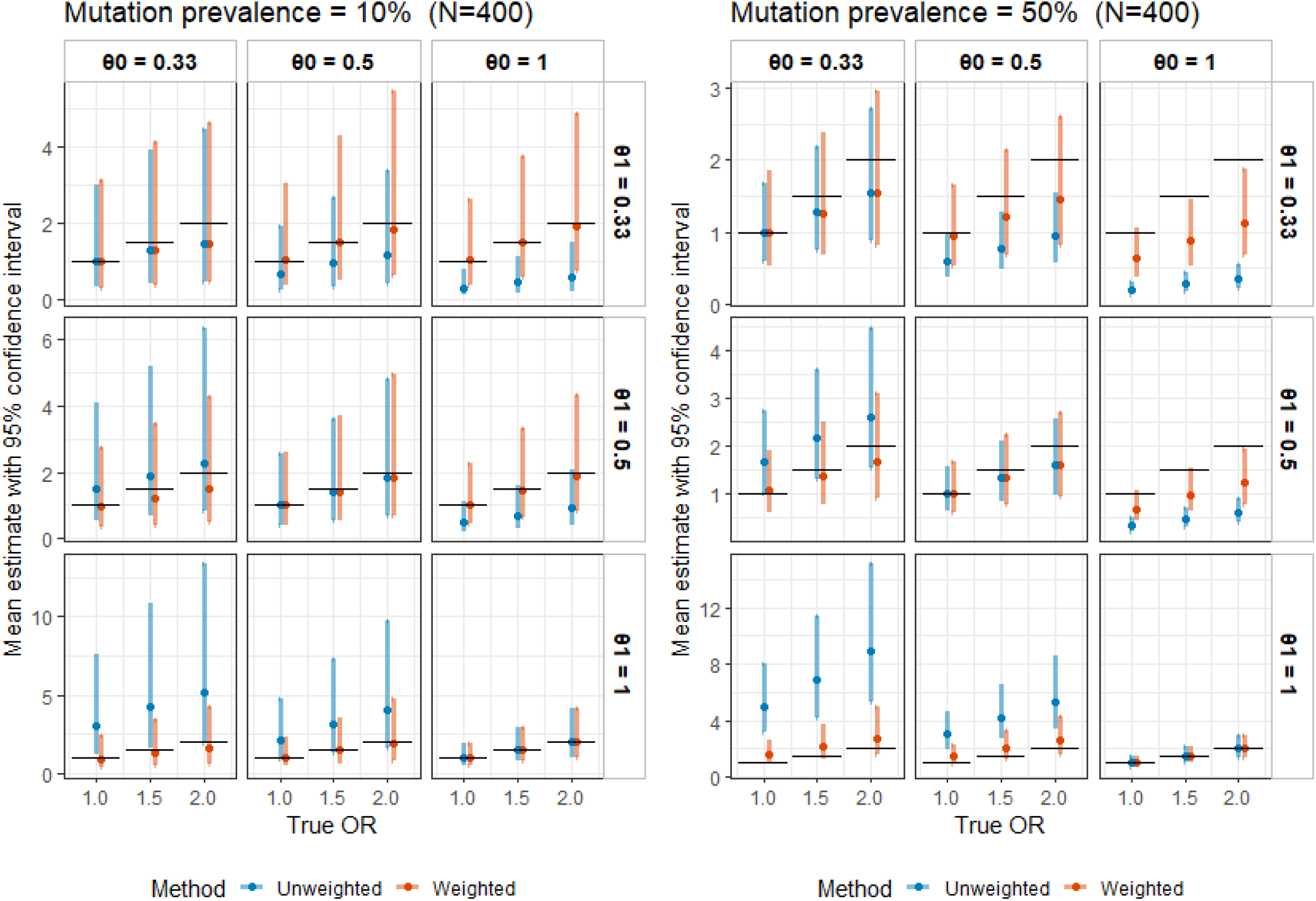
Simulation results: estimated OR with 95% confidence intervals under different settings (rows represent *θ*_1_, columns represent *θ*_0_); black bars represent the true OR).

#### Power and type I error of the comparison

The difference between the groups is based on the estimate of the OR, and tested using a Wald test to determine whether the OR is different from 1. A p-value of 0.05 or less indicates a significant difference. Power is defined as the proportion of significant tests under the alternative hypothesis (OR = 1.5 and OR = 2.0), while type I error is defined as the proportion of significant tests under the null hypothesis (OR = 1). Due to the large biases in the OR estimates, the unweighted method has a very large type I error rate in multiple scenarios, which means an inflation of false positive findings (Figure 3). On the other hand, the weighted approach better controls the type I error in general, although it is still high in some scenarios. The maximum type I error rate is 38.5% for the weighted approach, but reaches 100% for the unweighted approach. As a consequence, the power appears often higher with the unweighted method. Still, the power is acceptable for the weighted method, increasing reasonably as the effect size increases, except in scenarios where very few mutations are observed (mutation prevalence = 10%, *θ*_0_ = 0.33), which is expected.

**Figure 3.**
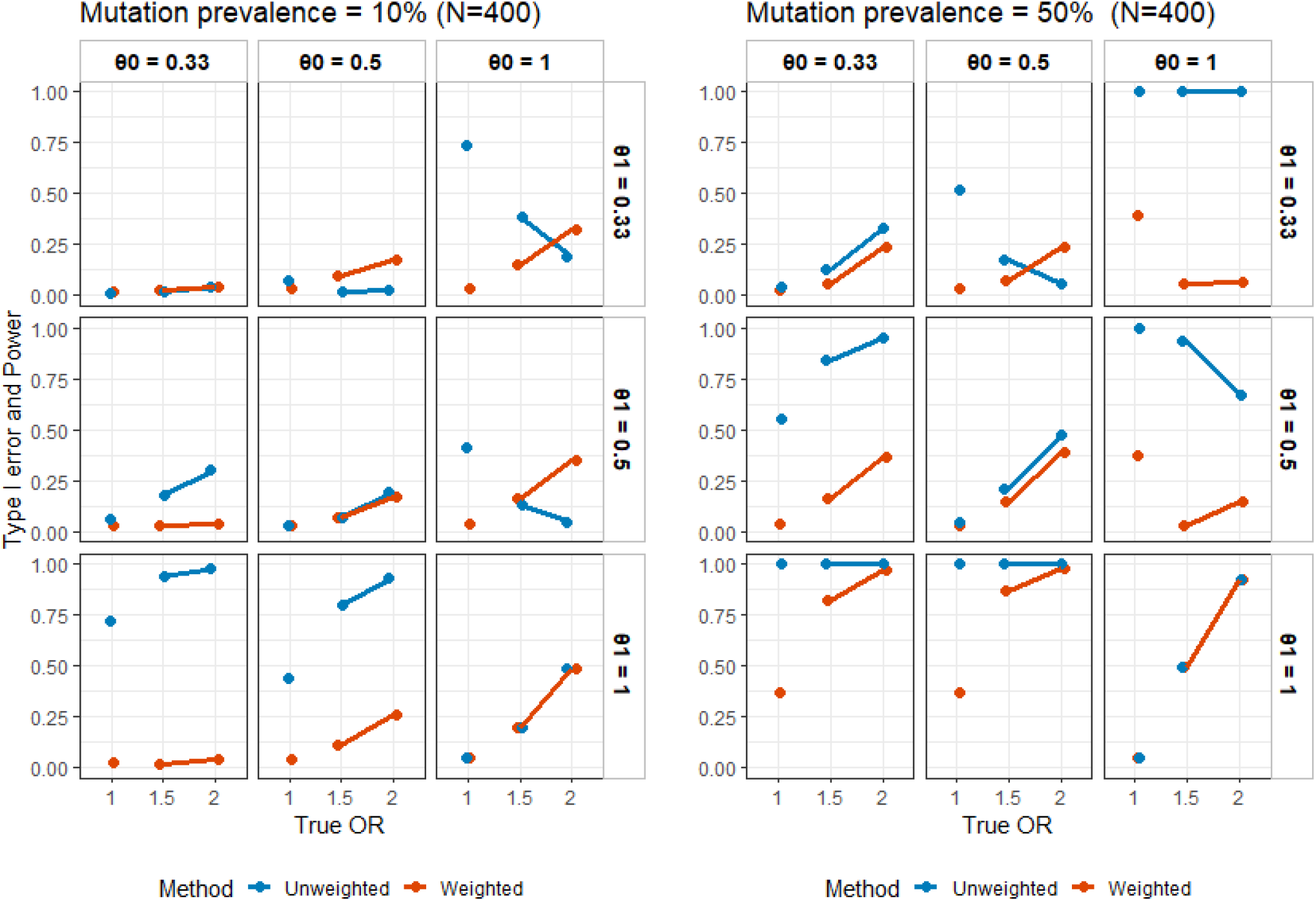
Simulation results: power of rejecting the null hypothesis OR=1 under different settings (rows represent *θ*_1_, columns represent *θ*_0_). Type I error and power are given in percentage.

All simulations were repeated with n=200 and n=1000 patients, and the results are presented in Appendix. The main conclusions were similar, although the number of scenarios where the true value was excluded from the confidence intervals was increased with n=1000 due to higher precision.

## Application

In many clinial studies, mutation call are often treated as binary variables, present or absent, without adjustment for potential uncertainty characterizing this call. Here we consider some examples where unequal mutation sensitivity in two groups is being suspected. For the purpose of our application, we assume they are no missing mutation calls due to technical errors in sequencing, insufficient sequencing depth, or bioinformatics pipeline. In the examples below, we propose using the average rescaled VAF of called mutations to adjust the estimate of the association between cohort and mutation as illustrated in Figure 1. Briefly, we assign each patient a weight, equal to 1 if mutation is observed, or unconditional average rescaled VAF (or potentially other metric of detection sensitivity) estimated from observed mutations within patient’s cohort. Inference can further be done using these weights applied within logistic regression or other likelihood based models.

### TCGA Kidney

Several genes are known to be important in kidney cancer. Previous study by Hakimi et al. (2013) explored tumor suppressors PBRM1, SETD2, and BAP1 for associations with cancer specific survival and tumor characteristics. Here, we illustrate the problem using a binary outcome, AJCC (American Joint Committee on Cancer) pathological stage. The TCGA Kidney cohort comprises of 356 patients with renal clear cell carcinoma that have tumor whole exome sequencing. Based on this cohort, we estimate the association between the presence of a mutation on three main genes and the stage as binary, defined as high (stage 3,4) vs low (stage 1,2), using a logistic regression with Wald tests. We found that somatic mutations in SETD2 and BAP1 are significantly associated with the higher tumor stage, with odds ratio (OR)=2.51 (p=0.003) and OR=3.38 (p=0.0006), respectively, while PBRM1 is not (OR=0.93, p=0.76). Among patients with high stage, 13% are SETD2 mutants, while among patients with low stage, only 7% are. As one can see in Figure 4, low stage patients with a mutation also have lower VAF, which potentially would mean there are more missed mutations in lower stage patients compared to high stage patients. Thus, lower mutation rate and lower stage may be confounded by higher rate of false negatives.

**Figure 4.**
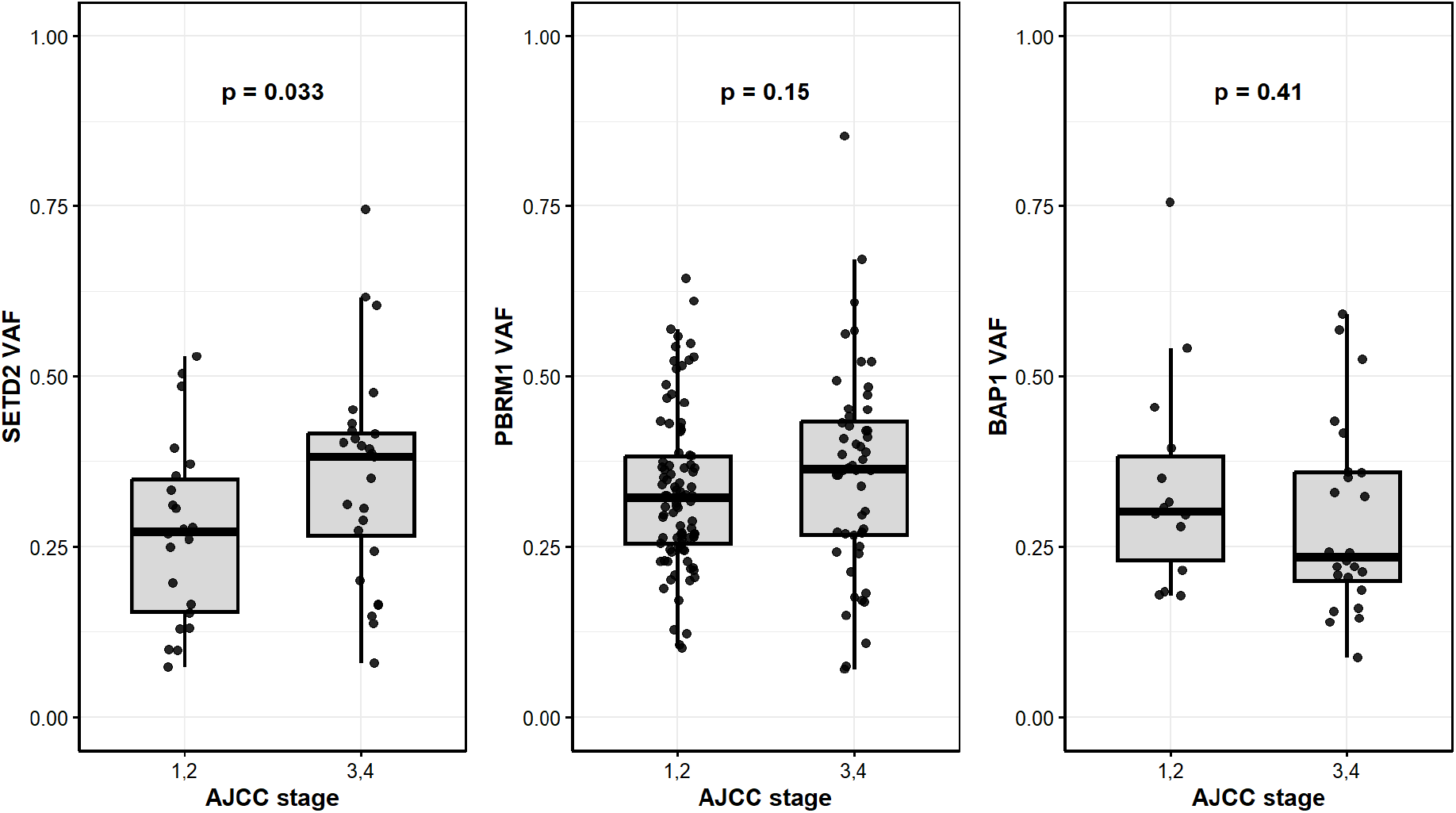
SETD2, PBRM1 and BAP1 VAF (variant allele frequency) by AJCC stage, p-value from Wilcoxon rank sum test

The adjusted ORs and p-values from our proposed weighted test become OR=1.8 (p=0.07), OR=4.17 (p=0.00015), and OR=0.97 (p=0.92) for SETD2, BAP1 and PBRM1, respectively. Thus, SETD2 is no longer significant.

### Immunotherapy response

Using a second published cohort form Gajic et al. (2022), another example is predicting response to immunotherapy from a sequenced pre-treatment tumor sample. If a patient is unlikely to respond they could be directed to another therapy. Traditionally, tumor mutational burden (TMB) serves as a biomarker of response, but individual genes were also implicated(Gajic et al. (2022)). This published cohort consists of six aggregated studies for a total of 319 patients with melanoma, non-small cell lung cancer, bladder cancer, and head and neck cancer, having available pre-immunotherapy whole exome sequencing. Among these, 272 were classified as responders or non-responders to immunotherapy (excluding stable disease). In that article, four genes, BCLAF1, TP53, KRAS and BRAF were found to be associated with treatment response in a multivariable model adjusted for TMB and age, tumor type and study of origin. Here we consider univariate associations of these genes, which might differ from published multivariable results with response for illustrative purpose: BCLAF1 is associated with non-response with OR 2.6 (p=0.04),TP53 has OR 0.9 (p=0.7), KRAS has OR 0.17 (p=0.003) and BRAF has OR 0.72 (p=0.26).

After incorporating adjustment for the VAFs, the associations become: BCLAF1 with OR 2.05 (p=0.19), TP53 has OR 0.61 (p=0.13), KRAS has OR 0.26 (p=0.029) and BRAF has OR 0.54 (p=0.094).

BRAF is an example where there was no significant association between mutation and response prior to adjustment, with 34% (32/94) of responders having a mutation on this gene vs 27% (48/178) of non-responders. However, as we can see from Figure 5, responders have significantly lower VAF, and potentially mutations are more likely to be missed among responders, and therefore 34% might be underestimated. This association is also supported by a study of immunotherapy in melanoma (van Not et al. (2022)).

**Figure 5.**
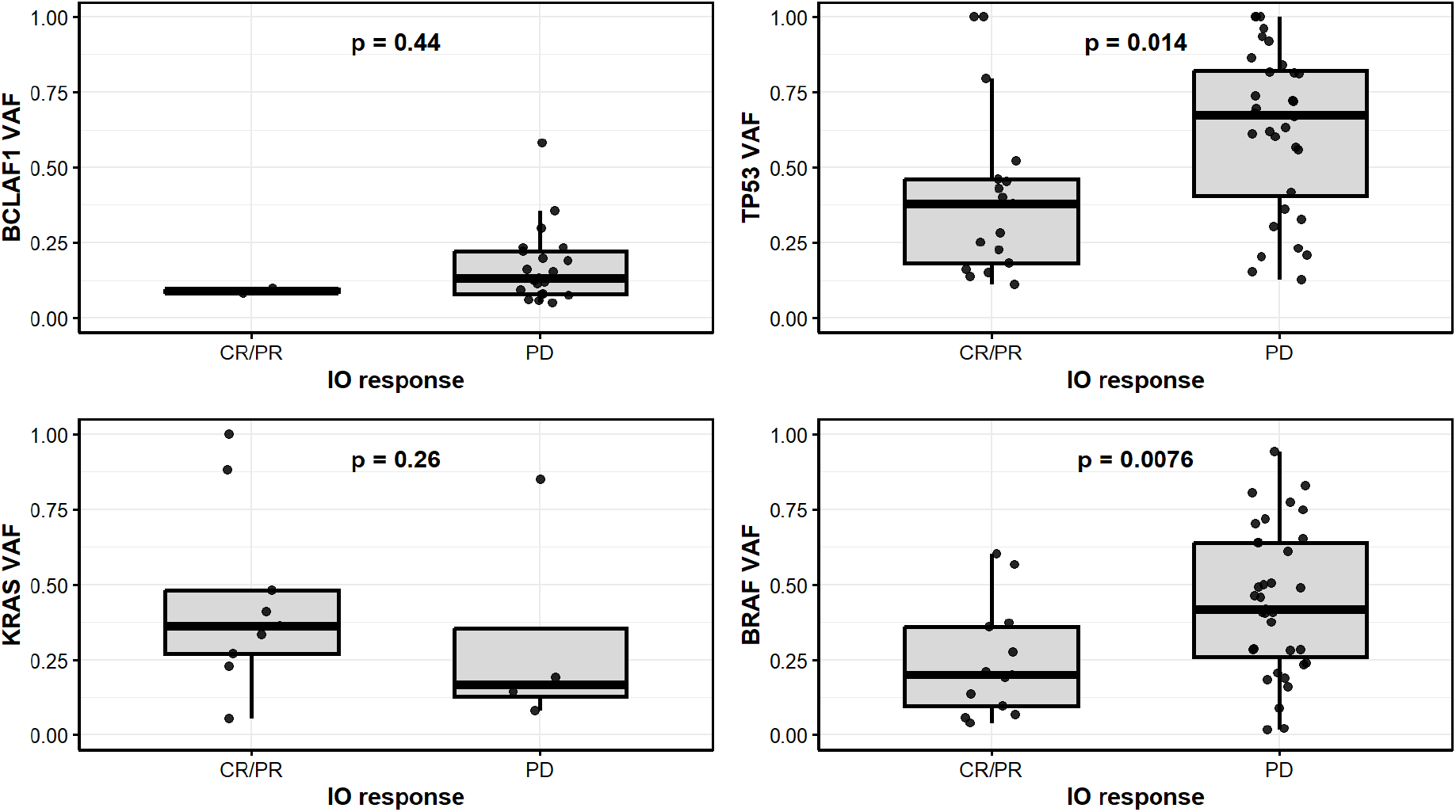
VAF for 4 genes associated with immunotherapy response. Percent of mutated patients is shown in each group, p-value from Wilcoxon rank sum test

For KRAS 12% (11/94) of responders had a mutation on this gene vs 2% (4/178) of non-responders. Although not significant, non-responders have lower VAF and potentially more missed mutations, suggesting that the true rates might be close.

### Bootstrap procedure

Surrogates for detection sensitivity such as VAF can be influenced by tumor purity, sequencing depth, and pipeline-specific variant calling decisions, making it an unstable proxy for true mutation prevalence. To reduce sensitivity to the arbitrary overall normalization of the weights, we propose to use a bootstrap procedure to estimate confidence intervals. This procedure also helps to account for uncertainty arising from estimation of the group-specific detection sensitivities. Subjects are resampled with replacement, 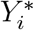, within outcome groups if appropriate, and the weights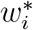 are recalculated based on 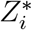, and before refitting the weighted logistic regression. Confidence intervals for 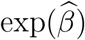 are obtained using percentiles from the bootstrap distribution. The confidence interval can be estimated using percentiles of bootstrapped ORs.

In the examples we considered, in TCGA data, SETD2 changed from unadjusted OR 2.5 (95% CI 1.4, 4.6) to weighted OR and bootstrap CI of OR 1.8 (95% CI 0.9, 3.7), BAP1 went from OR 3.4 (95% CI 1.7, 6.8) to OR 4.2 (95% CI 2.0, 9.9), and PBRM1 from OR 0.93 (95% CI 0.6, 1.4) to 0.98 (95% CI 0.6,1.6). In the immunotherapy dataset, BCLAF1 changed from unadjusted OR 2.6 (95% CI 1.04, 6.6) to weighted OR and bootstrap CI of 2.05 (95% CI 0.01, 5.4), TP53 went from OR 0.9 (95% CI 0.5, 1.6) to OR 0.6 (95% CI 0.3, 1.1), KRAS from OR 0.2 (95% CI 0.05, 0.6) to OR 0.26 (95% CI 0.03, 1.3) and BRAF from OR 0.7 (95% CI 0.4, 1.2) to OR 0.5 (95% CI 0.2, 1.5).

Thus, in the TCGA kidney cancer data, adjustment for mutation detectability had heterogeneous effects across genes. For SETD2, evidence was inconclusive after adjustment. In contrast, the BAP1 association was strengthened, and for PBRM1, both the point estimate and confidence interval changed very little. In the immunotherapy dataset, weighting generally altered the magnitude of the estimated associations while the bootstrap confidence intervals became substantially wider. The BCLAF1 estimate was attenuated while the weighted estimates for TP53, KRAS, and BRAF remained in broadly the same direction as their unadjusted estimates with bootstrap 95% CI including 1.

These results illustrate that adjustment for differential mutation detectability can affect both the magnitude and direction of estimated associations, while bootstrap inference accounting for uncertainty in the estimated weights may substantially reduce precision. Thus, the method should not be viewed as systematically strengthening associations, but as correcting point estimates while appropriately reflecting the additional uncertainty introduced by estimating mutation detectability.

## Discussion

A question universally assessed in all types of cancer is what mutation mechanism is at play. Identifying which specific mutations are associated with different tumor characteristics or response to certain therapies can further the understanding of the disease and guide patients’ care, ultimately leading to improved outcomes. However, mutations can sometimes be missed due to tumor heterogeneity, making it difficult to make a reliable estimation of such associations. In our simulation study, it was confirmed that missed mutations impact the type I error and power of tests used to define these associations, especially when one of the compared group has a higher rate of missed mutations than the other. The highest impact was seen on the type I error, which was not controlled when the group with the lower mutation prevalence also had the higher risk of missed mutation, leading to underestimation of the mutation prevalence in that group, and overestimation of the difference between the groups. In such cases, the estimated OR far exceeded the true one, with confidence interval boundaries excluding the true OR. To address the issue, we proposed the use of a weighted estimate to correct for the risk of missing a mutation when estimating the mutation prevalence and comparing it between two groups. The weight used is based on the available data, and should reflect as much as possible the absolute risk of missing a mutation. Our study found that the use of a weighted model reduced the bias in the OR estimate, leading to confidence intervals covering the true OR in almost all cases. This bias correction came at the price of lower power but better controlled type I error.

Bias from misclassification in binary regression models is well documented, and can be severe unless classification accuracy is very high (Neuhaus (1999)), motivating explicit correction methods in measurement-error frameworks (Carroll et al. (2006)). Our contribution addresses a distinctive genomics feature: the detection probability is individual-level and its surrogate (e.g., VAF) is observed only when a mutation is called, so the sensitivity must be inferred from the observed conditional distribution.

The presence of intra-tumor heterogeneity is a well accepted issue that results in potentially capturing only a subset of tumor’s mutational profile (McGranahan and Swanton (2017)). Such missed mutations can obscure the true clonal architecture of the tumor and underestimate the complexity of evolutionary dynamics, which in turn hampers the identification of driver events and therapeutic targets (Turajlic et al. (2019)). Some technical ways to handle it include increasing the sequencing depth to maximize the chance of picking subclonal mutation, and including multiple callers in a pipeline (Callari et al. (2017)), although it does not address the issue of spatial heterogeneity. Multiregional sequencing Gerlinger et al. (2014) is shown to improve the accuracy of mutation detection, as well as combining multiple tumor cores (pooling), prior to sequencing (DiNatale et al. (2021), Pongor et al. (2020)), but has not been widely adopted due to added costs and labor. In addition, liquid biopsy using ctDNA might supplement tissue biopsy in order to make mutation detection more accurate (Ma et al. (2024)).

We assume that there are no false positives, i.e. if mutation is detected, it is in a sequenced sample. Thus, we ignore technical artefacts, i.e. mutation is called but it’s not there. In support of these assumption is increasing quality of sequencing technology and calling algorithms. On the other hand, false negatives that are due to tumor heterogeneity will not get better with improvement in bulk sequencing technology.

The main pitfall of the proposed approach is that it relies on weights that has to be estimated from the data. Ideally, they should be calibrated to reflect the actual probability that mutation is missed, given that the mutation is there. In reality, this cannot be directly measured or known. We have to rely on surrogate metrics, such as the VAF. By representing how many of the reads have the variant allele, it shows how likely it is to find this variant. However, it is not available in subjects for whom the mutation was not called, and the weights for these have to be extrapolated based on the rescaled VAF in subjects with mutations. Thus, we are making an assumption that the detection sensitivity in general, or in this case VAFs, come from the some probability distribution, potentially outcome specific. This allows for patient variability in detection sensitivity. Other alternatives could be used, such as the cancer cell fraction (CCF), but it is not widely available for published datasets. Further work is needed to help identify the best weight that is both available and meaningfully represent the desired probability. Explorations of large datasets of multiregional sequencing could help compare different weights definitions.

One advantage of the weights being derived directly within the cohort of interest and not relying on outside information, as that the weights will likely be based on patients with similar diseases, similar therapies, and similar sequencing platforms, at least similar enough that they are part of a unique study. It increases the applicability of the proposed method to a diverse range of studies. However, due to the limited nature of the illustrative application, the current scope of application is mainly for solid tumors in adults, in studies where tumor heterogeneity is expected. We see the scope of applicability is whenever tumor heterogeneity in a certain gene is substantial, especially if it is differential between the two groups. The second, major advantage of proposing a weighted approach is its versatility. It could be used in regression context, such as the logistic regression presented here, or in the context of a z-test. Weights can be adapted for other types of outcomes, e.g. linear or time-to-event outcomes, using parametric model. This makes the proposed approach easily applicable in practice, notably with the R code that can be accessed at https://github.com/IOstrovnaya/mutation-sensitivity-weights/tree/main.

## Conclusions

This study confirmed that, in patients with cancer, missed mutations due to tumor heterogeneity can be problematic when evaluating the association of a given mutation with binary outcome. Using a weighted approach to correct for the risk of a mutation to be missed helps mitigate the inflated type I error often seen in such situations.

## Availability of data and code

The data used in this manuscript can be publicly accessed through https://www.cbioportal.org/study/summary?id=kirc_tcga_pan_can_atlas_2018 and https://gitlab.com/sanjanalab/circle. The R code to conduct the proposed weighted comparisons is available on github at https://github.com/IOstrovnaya/mutation-sensitivity-weights.

## Authors’ Contributions

AM and IO contributed equally to the design of the study, the analysis of the data and writing of the manuscript.

## Appendix

**Simulation results with n=200**

**Simulation results with n=1000**

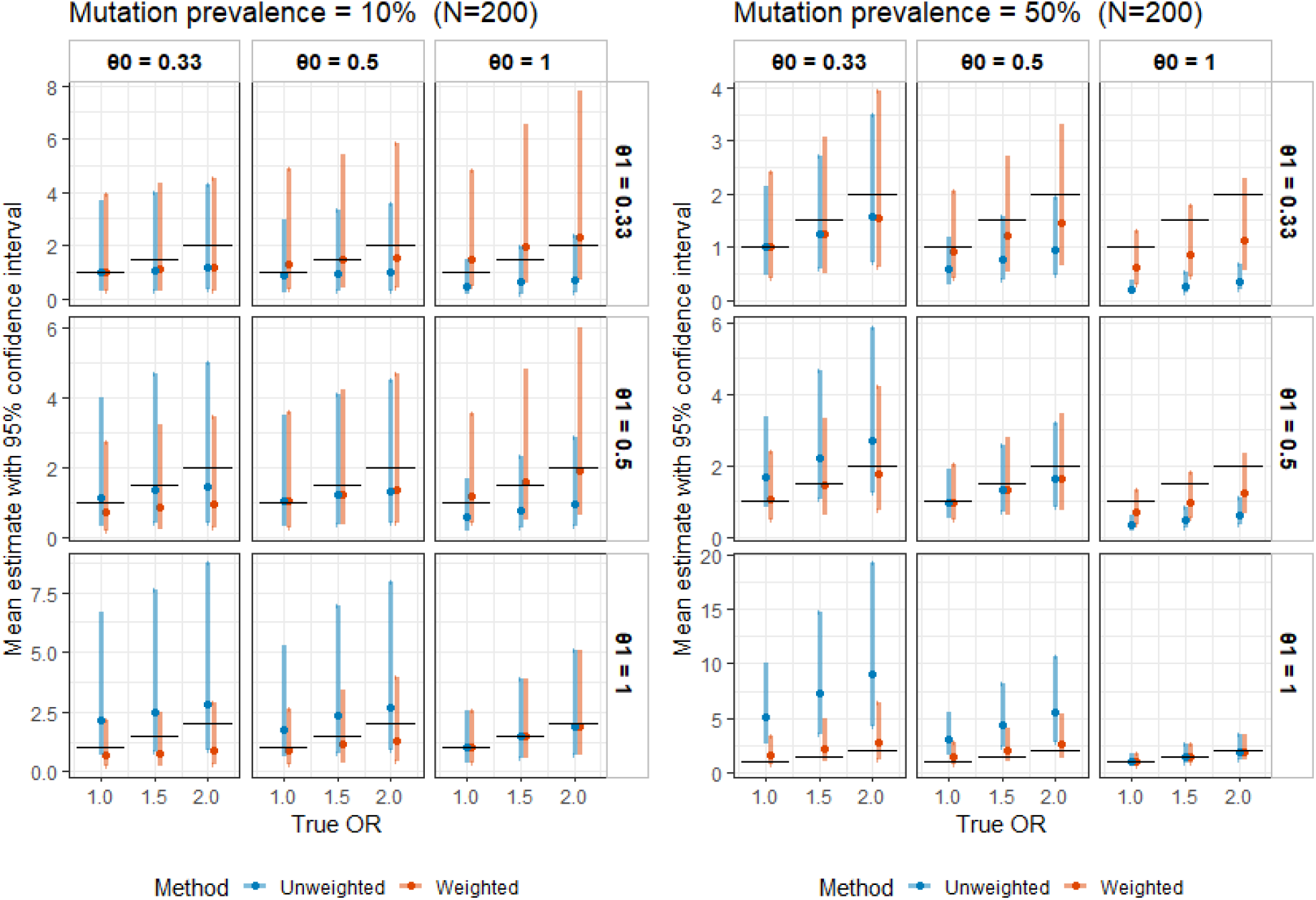

Simulation results: estimated OR with 95% confidence intervals under different settings (rows represent *θ*_1_, columns represent *θ*_0_); black bars represent the true OR).

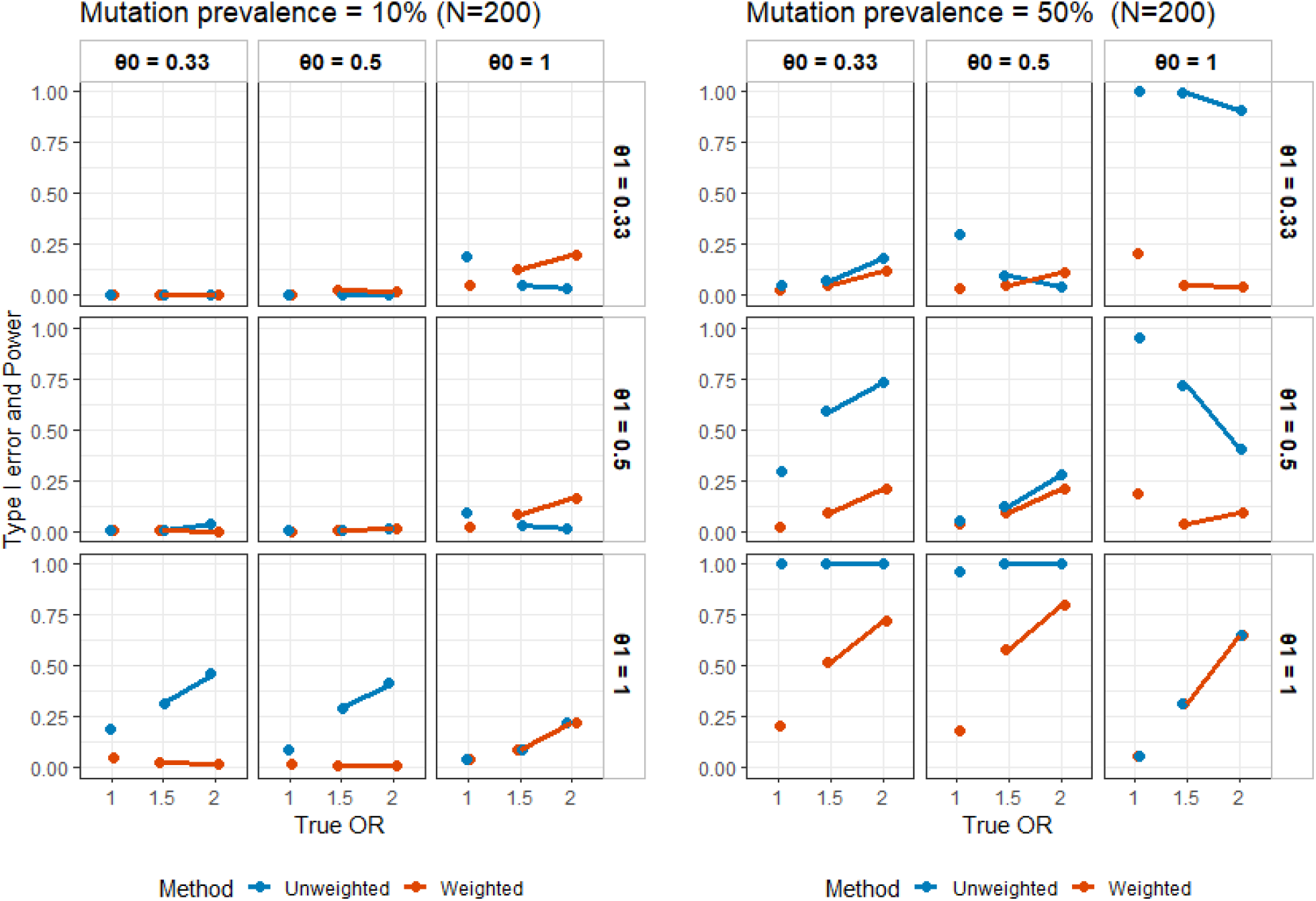

Simulation results: power of rejecting the null hypothesis OR=1 under different settings (rows represent *θ*_1_, columns represent *θ*_0_). Type I error and power are given in percentage.

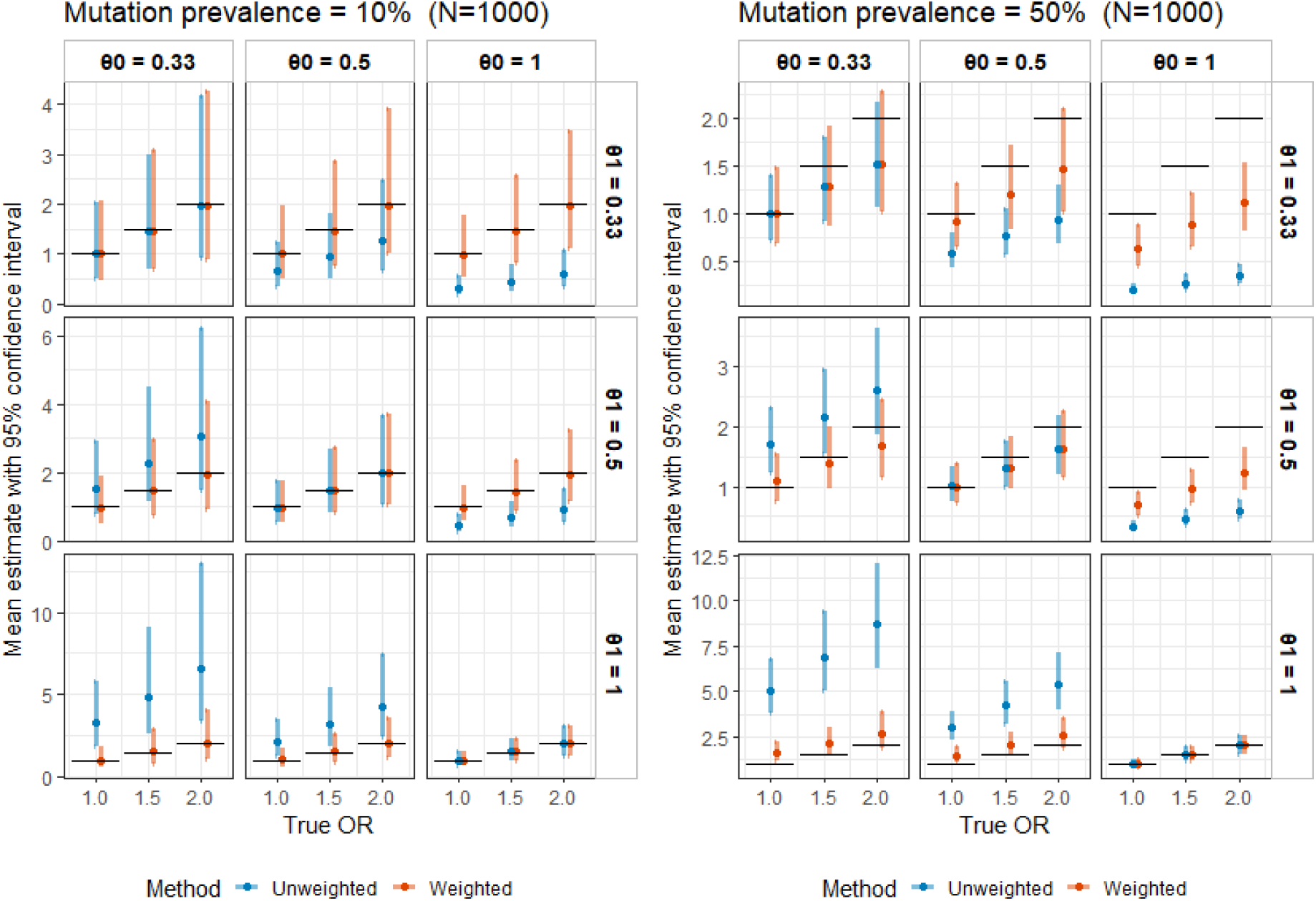

Simulation results: estimated OR with 95% confidence intervals under different settings (rows represent *θ*_1_, columns represent *θ*_0_); black bars represent the true OR).

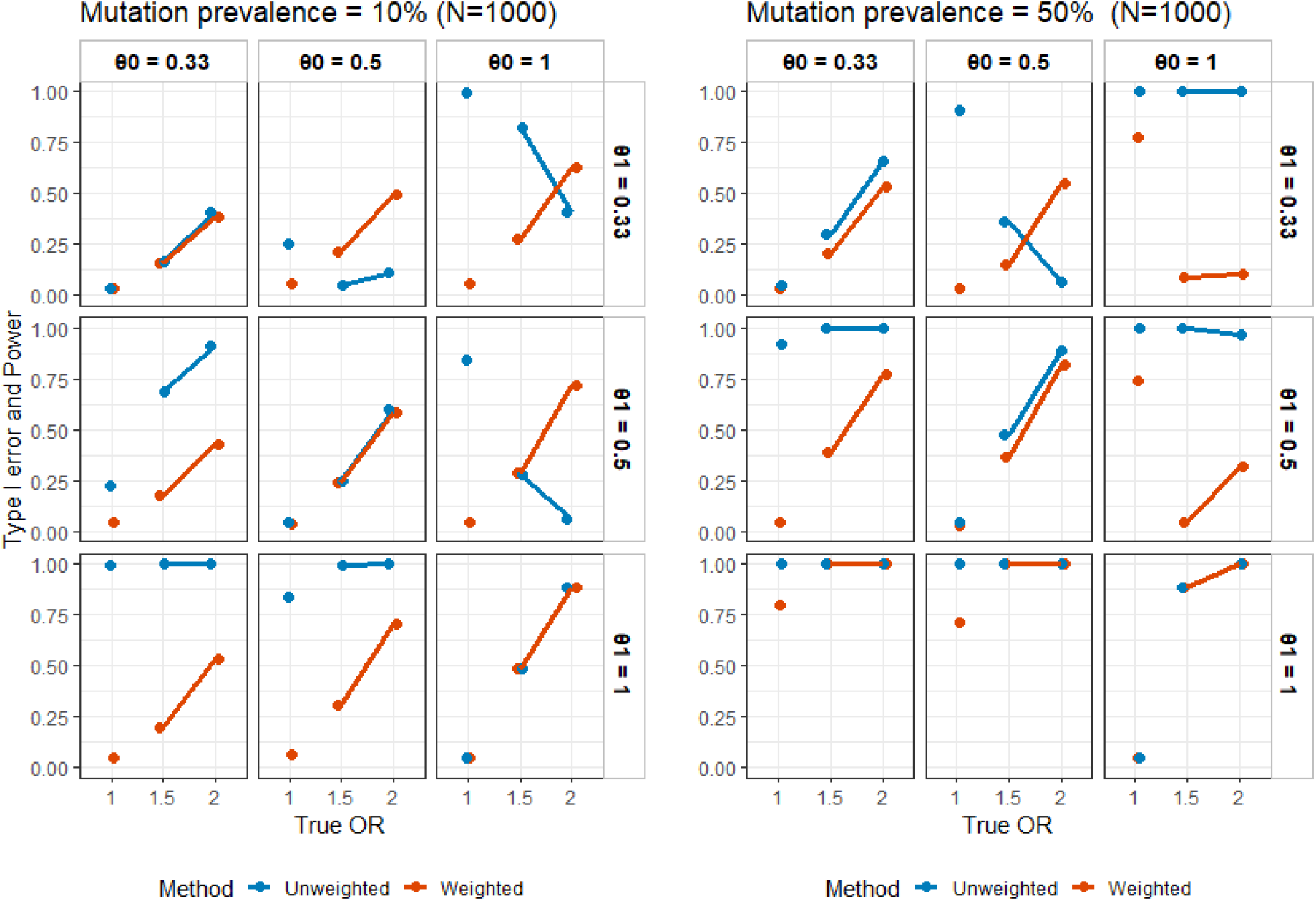

Simulation results: power of rejecting the null hypothesis OR=1 under different settings (rows represent *θ*_1_, columns represent *θ*_0_). Type I error and power are given in percentage.

## References

Callari, M., Sammut, S.-J., De Mattos-Arruda, L., Bruna, A., Rueda, O. M., Chin, S.-F., and Caldas, C. (2017). Intersect-then-combine approach: improving the performance of somatic variant calling in whole exome sequencing data using multiple aligners and callers. Genome Medicine, 9(1):35.

Carroll, R. J., Ruppert, D., Stefanski, L. A., and Crainiceanu, C. M. (2006). Measurement Error in Nonlinear Models. Chapman and Hall/CRC, 0 edition.

DiNatale, R. G., Mano, R., Makarov, V., Rusk, N., Drill, E., Winer, A., Sankin, A., Yoo, A., Freeman, B. A., Hsieh, J. J., Chen, Y.-B., Coleman, J. A., Berger, M., Ostrov-naya, I., Chan, T. A., Russo, P., Reznik, E., and Hakimi, A. A. (2021). Resource-efficient pooled sequencing expands translational impact in solid tumors. Kidney Cancer Journal: Official Journal of the Kidney Cancer Association, 19(2):18–23.

Gajic, Z. Z., Deshpande, A., Legut, M., Imieliński, M., and Sanjana, N. E. (2022). Recurrent somatic mutations as predictors of immunotherapy response. Nature Communications, 13(1):3938.

Gerlinger, M., Horswell, S., Larkin, J., Rowan, A. J., Salm, M. P., Varela, I., Fisher, R., McGranahan, N., Matthews, N., Santos, C. R., Martinez, P., Phillimore, B., Begum, S., Rabinowitz, A., Spencer-Dene, B., Gulati, S., Bates, P. A., Stamp, G., Pickering, L., Gore, M., Nicol, D. L., Hazell, S., Futreal, P. A., Stewart, A., and Swanton, C. (2014). Genomic architecture and evolution of clear cell renal cell carcinomas defined by multiregion sequencing. Nature Genetics, 46(3):225–233.

Hakimi, A. A., Ostrovnaya, I., Reva, B., Schultz, N., Chen, Y.-B., Gonen, M., Liu, H., Takeda, S., Voss, M. H., Tickoo, S. K., Reuter, V. E., Russo, P., Cheng, E. H., Sander, C., Motzer, R. J., Hsieh, J. J., and ccRCC Cancer Genome Atlas (KIRC TCGA) Research Network investigators (2013). Adverse outcomes in clear cell renal cell carcinoma with mutations of 3p21 epigenetic regulators BAP1 and SETD2: a report by MSKCC and the KIRC TCGA research network. Clinical Cancer Research: An Official Journal of the American Association for Cancer Research, 19(12):3259–3267.

Jamal-Hanjani, M., Wilson, G. A., McGranahan, N., Birkbak, N. J., Watkins, T. B. K., Veeriah, S., Shafi, S., Johnson, D. H., Mitter, R., Rosenthal, R., Salm, M., Horswell, S., Escudero, M., Matthews, N., Rowan, A., Chambers, T., Moore, D. A., Turajlic, S., Xu, H., Lee, S.-M., Forster, M. D., Ahmad, T., Hiley, C. T., Abbosh, C., Falzon, M., Borg, E., Marafioti, T., Lawrence, D., Hayward, M., Kolvekar, S., Panagiotopoulos, N., Janes, S. M., Thakrar, R., Ahmed, A., Blackhall, F., Summers, Y., Shah, R., Joseph, L., Quinn, A. M., Crosbie, P. A., Naidu, B., Middleton, G., Langman, G., Trotter, S., Nicolson, M., Remmen, H., Kerr, K., Chetty, M., Gomersall, L., Fennell, D. A., Nakas, A., Rathinam, S., Anand, G., Khan, S., Russell, P., Ezhil, V., Ismail, B., Irvin-Sellers, M., Prakash, V., Lester, J. F., Kornaszewska, M., Attanoos, R., Adams, H., Davies, H., Dentro, S., Taniere, P., O’Sullivan, B., Lowe, H. L., Hartley, J. A., Iles, N., Bell, H., Ngai, Y., Shaw, J. A., Herrero, J., Szallasi, Z., Schwarz, R. F., Stewart, A., Quezada, S. A., Quesne, J. L., Loo, P. V., Dive, C., Hackshaw, A., and Swanton, C. (2017). Tracking the Evolution of Non–Small-Cell Lung Cancer. New England Journal of Medicine, 376(22):2109–2121.

Landau, D. A., Carter, S. L., Stojanov, P., McKenna, A., Stevenson, K., Lawrence, M. S., Sougnez, C., Stewart, C., Sivachenko, A., Wang, L., Wan, Y., Zhang, W., Shukla, S. A., Vartanov, A., Fernandes, S. M., Saksena, G., Cibulskis, K., Tesar, B., Gabriel, S., Hacohen, N., Meyerson, M., Lander, E. S., Neuberg, D., Brown, J. R., Getz, G., and Wu, C. J. (2013). Evolution and impact of subclonal mutations in chronic lymphocytic leukemia. Cell, 152(4):714–726.

Liu, C., Liu, A., Hu, J., Yuan, V., and Halabi, S. (2014). Adjusting for misclassification in a stratified biomarker clinical trial. Statistics in Medicine, 33(18):3100–3113.

Ma, L., Guo, H., Zhao, Y., Liu, Z., Wang, C., Bu, J., Sun, T., and Wei, J. (2024). Liquid biopsy in cancer: current status, challenges and future prospects. Signal Transduction and Targeted Therapy, 9(1):1–36.

Magder, L. S. and Hughes, J. P. (1997). Logistic Regression When the Outcome Is Measured with Uncertainty. American Journal of Epidemiology, 146(2):195–203.

McGranahan, N. and Swanton, C. (2017). Clonal Heterogeneity and Tumor Evolution: Past, Present, and the Future. Cell, 168(4):613–628.

Morris, L. G., Riaz, N., Desrichard, A., şenbabaoğlu, Y., Hakimi, A. A., Makarov, V., Reis-Filho, J. S., and Chan, T. A. (2016). Pan-cancer analysis of intratumor heterogeneity as a prognostic determinant of survival. Oncotarget, 7(9):10051–10063.

Neuhaus, J. (1999). Bias and efficiency loss due to misclassified responses in binary regression. Biometrika, 86(4):843–855.

Pongor, L. S., Munkácsy, G., Vereczkey, I., Pete, I., and Győrffy, B. (2020). Currently favored sampling practices for tumor sequencing can produce optimal results in the clinical setting. Scientific Reports, 10(1):14403.

Turajlic, S., Sottoriva, A., Graham, T., and Swanton, C. (2019). Resolving genetic heterogeneity in cancer. Nature Reviews Genetics, 20(7):404–416.

van Not, O. J., Blokx, W. A. M., van den Eertwegh, A. J. M., de Meza, M. M., Haanen, J. B., Blank, C. U., Aarts, M. J. B., van den Berkmortel, F. W. P. J., de Groot, J. W. B., Hospers, G. A. P., Kapiteijn, E., Piersma, D., van Rijn, R. S., Stevense-den Boer, M., van der Veldt, A. A. M., Boers-Sonderen, M. J., Jansen, A. M. L., Wouters, M. W. J. M., and Suijkerbuijk, K. P. M. (2022). BRAF and NRAS Mutation Status and Response to Checkpoint Inhibition in Advanced Melanoma. JCO precision oncology, 6:e2200018.

Vitale, I., Shema, E., Loi, S., and Galluzzi, L. (2021). Intratumoral heterogeneity in cancer progression and response to immunotherapy. Nature Medicine, 27(2):212–224. For data citations of datasets uploaded to e.g. figshare, please use the howpublished option in the bib entry to specify the platform and the link, as in the Hao:gidmaps:2014 example in the sample bibliography file.

